# High-speed atomic force microscopy reveals surface-catalyzed elongation mechanism of fungal functional amyloid, hydrophobin RolA

**DOI:** 10.1101/2025.08.19.671040

**Authors:** Nao Takahashi, Tatsuya Kimura, Yuki Terauchi, Takumi Tanaka, Akira Yoshimi, Takahiro Watanabe-Nakayama, Keietsu Abe

**Author notes:** **Author Contributions:** N.T., T.W.-N., and K.A. designed the research; N.T., K.T., and T.W.-N. performed the research; N.T., and T.W.-N. analyzed the data; N.T., T.W.-N., Y.T., T.T., A.Y., and K.A. wrote the paper; All authors discussed the results and commented on the manuscript; All authors have approved the final version of the manuscript; K.A. supervised this research. **Competing Interest Statement:** The authors declare no competing interest.

## Abstract

Hydrophobins, a type of functional amyloid, are conserved in filamentous fungi and act as a protective coat in the fibrous form called rodlet. Rodlets form hierarchical structures where they are bundled and densely aligned, contributing to the hydrophobicity of the mycelium surface. However, the formation mechanism of the hierarchical structures is completely unknown. In this study we used high-speed atomic force microscopy to directly observe the structural dynamics of hierarchical structure formation by hydrophobin RolA from the industrial fungus *Aspergillus oryzae* at a single-fibril level and revealed its mechanism. The elongation of rodlets occurred at both ends and was discontinuous, alternating between periods when they could elongate (growth state) and could not elongate (pause state). This suggests an equilibrium of two distinct structural states at the rodlet ends. We also discovered an aggregation pathway, termed “surface-catalyzed elongation”, in which elongation is promoted by lateral interactions between bundled rodlets. Surface-catalyzed elongation decreased the energy barrier of both structural switching between growth and pause states and elongation at rodlet ends, doubling the elongation rate in bundled rodlets. The rodlet surface could be considered as a catalyst for the elongation of neighboring rodlets. Surface-catalyzed elongation could contribute to rodlet bundling, whereby rodlets tend to form oriented domain structures, and our Monte Carlo simulations confirmed this. Surface-catalyzed elongation may be a universal concept to explain the hierarchical assembly mechanism of amyloid fibrils, so it could contribute to the advancement of amyloid research in general.

## Introduction

Assemblies of amyloid proteins into cross-β fibrils, which could form further huge aggregates with bundling (1, 2), are beneficially used as functional amyloids by bacteria, fungi, insects, and mammals to maintain physiological processes (3, 4), although amyloid fibrils are often associated with protein misfolding and neurodegenerative diseases. Hydrophobins are functional amyloids produced by filamentous fungi; they form amyloid-like fibrils called rodlets, which become ordered to form dense rodlet films and support fungal physiology (5, 6). Hydrophobins are low-molecular-weight amphiphilic proteins that adhere to the cell wall surface to coat and make it hydrophobic. This surface modification helps to promote air dispersibility of conidia and helps hyphae to adhere to hydrophobic surfaces such as plant leaves coated with wax esters (7, 8). Hydrophobins also have immunosilencing properties, because hydrophobin coating of the hyphae of pathogenic filamentous fungi prevents recognition by the host immune system (5, 9-11). Some hydrophobins may be involved in the degradation of solid polymers (12-15).

Hydrophobins are classified into several classes based on their amino acid sequences and surface hydrophobicity. Class I hydrophobins self-assemble into rodlets. Hydrophobins typically contain three hydrophobic loops that form cross β-sheet structures, which assemble into bundled ordered rodlets on the conidial surface (5, 6, 16). In our previous studies, purified hydrophobin RolA from the industrial fungus *Aspergillus oryzae* formed dense rodlet films at the air–water interface and the films were transferred to the substrate, significantly altering substrate wettability (17, 18). Therefore, rodlets are key factors that affect solid surface properties and may regulate biological functions of the fungal cell wall, although the mechanisms of their formation remain largely unknown. Understanding the dynamics of rodlet film formation, where hydrophobins transition from monomers to rodlets, which then assemble into dense hierarchical structures, is key to uncovering the diverse functions of hydrophobins.

Rodlet formation is thought to resemble usual amyloid fibril formation: it is initiated by nucleation, followed by autocatalytic growth as monomers bind to rodlet ends using the preformed rodlets as templates (14, 19). Previous studies that used thioflavin T (ThT) assays and atomic force microscopy (AFM) with coarse temporal resolution have provided macroscopic insights into rodlet film formation but failed to reveal the detailed mechanism (5, 16, 18). A comprehensive understanding of rodlet film formation requires elucidating not only the formation of individual rodlets but also their interactions and assembly. This necessitates high spatial and temporal resolution of real-time observations at the single-fibril level and quantitative analysis of local interactions.

In this study, we used high-speed (HS) AFM (20) to observe formation of the rodlets of hydrophobin RolA from *A. oryzae* and to directly measure the kinetics of rodlet formation at a single-fibril level. HS-AFM showed that rodlets elongate discontinuously and unveiled their remarkable ability to elongate rapidly from both ends. In addition, the kinetics of elongation depend on the presence of neighboring rodlets, with elongation accelerating in their vicinity. The lateral interactions of rodlets led to their bundling, and our Monte Carlo simulations showed that this local reaction, detected by HS-AFM, contributed to the collective ordering of the dense rodlet film. These findings provide new insights into the governing principles behind the hierarchical assembly of amyloid fibrils.

## Results

### Rodlet formation in bulk solution

To investigate structural changes in RolA associated with rodlet formation, we used circular dichroism (CD) spectroscopy. The CD spectrum of monomeric RolA had a minimum at approximately 205 nm (Figure S1*A*), consistent with our previous findings (13). Upon rodlet formation, the minimum shifted to approximately 230 nm (Figure S1*B*), indicating an increase in β-sheet content compared with that in the monomeric state (21). These results support the hypothesis that RolA undergoes rodlet formation through the establishment of intermolecular β-sheet structures.

To further elucidate the mechanism of rodlet formation, we performed a ThT fluorescence assay. The reaction curves did not have the characteristic sigmoidal shape typically observed in amyloid protein aggregation (19). Instead, an immediate increase in fluorescence intensity was observed without a lag phase, followed by a constant rate of fluorescence increase until saturation was reached (Figure S1*C*). Data fitting with Eq. 1 (see Methods) provided good fits (Figure S1*D*). Model fitting showed that rodlet formation was well described by the dock–lock model and the absence of a rodlet-replication system, such as secondary nucleation (14, 22, 23), as reported previously for other class I hydrophobins (14). However, the above analysis did not provide further mechanistic insight into rodlet formation, highlighting the limitations of bulk solution measurements in elucidating the details of this process.

### HS-AFM analysis of rodlet elongation

To elucidate the details of the mechanism of rodlet elongation, we examined it at the single-fibril level using HS-AFM. A small amount of rodlets was first immobilized on a hydrophobic silicon substrate, and then the substrate was immersed in a RolA monomer solution (7.35 µM) during the measurement. We chose a hydrophobic substrate for this analysis because RolA adsorbs to solid surfaces via hydrophobic interactions (24) and can form rodlets at hydrophobic–hydrophilic interfaces (17).

Measurements revealed frequent elongation of pre-existing rodlets, as well as the formation and subsequent elongation of newly generated rodlets (de novo rodlets) (Figure 1*A*, Supporting video 1). Cross-sectional analysis revealed that rodlets had an approximate height of 3 nm (Figure 1*B*). Spherical aggregates formed before rodlet emergence and served as rodlet precursors: their height gradually increased and reached approximately 3 nm, followed by a structural transformation into rod-like formations, which subsequently initiated elongation (Figure 1*C*).

**Figure 1.**
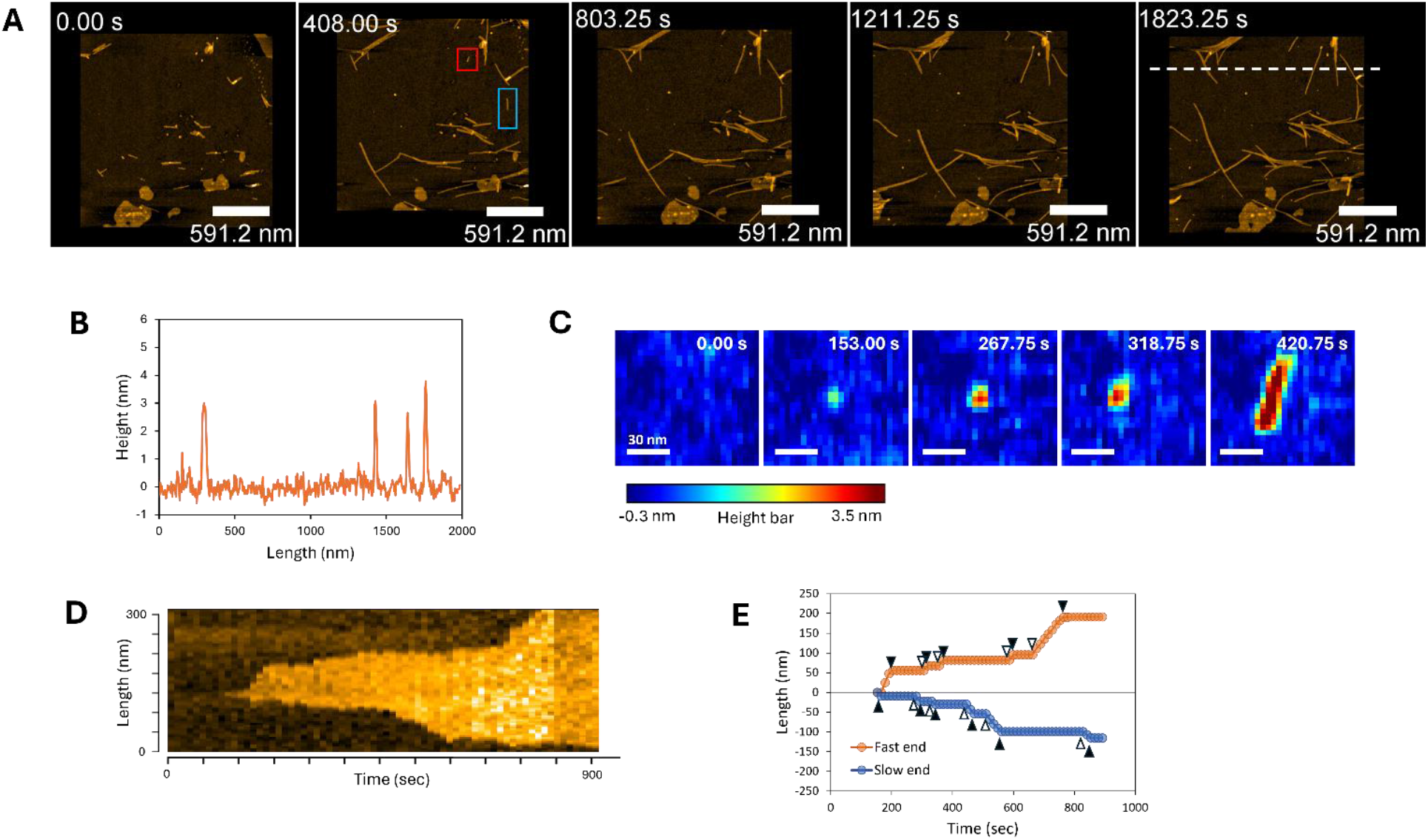
High-speed atomic force microscopy imaging of RolA rodlet formation. (A) Images taken during rodlet formation (scan size, 2 μm × 2 μm; pixels along the *X-*axis, 500; pixels along the *Y-*axis, 250). (B) Cross-section at the white dashed line in (A). (C) Representative images of the area within the red square in (A), showing that a spherical structure was formed first, followed by rodlet formation. (D) Representative kymograph of the area within the blue rectangle in (A), showing that the rodlet elongated from both ends in a stepwise manner. (E) Time evolution of the positions of rodlet ends, showing that both ends repeatedly grew and stopped. Black triangles, start points of dwell time; white triangles, start points of step time.

Pre-existing rodlets initially formed at the air–water interface, whereas de novo rodlets appeared on the substrate. Because amyloid fibril elongation can vary depending on the structural properties of the template surface (25), we compared the elongation behavior of pre-existing and de novo rodlets. HS-AFM video analysis revealed that rodlets alternated between periods of elongation and periods of length stabilization (Figure 1*D* and *E*). The apparent elongation rate, determined from the slope of a linear fit connecting the onset and termination points of rodlet elongation, was 7.9 ± 0.9 nm/min (mean ± S.E.) for pre-existing rodlets and 11.6 ± 2.4 nm/min for de novo rodlets, with no statistically significant difference (Figure S2*A*, Table *S1*).

Dwell time (duration of no length change), step time (duration of elongation), and step size (elongation length per step time) for both pre-existing and de novo rodlets followed a typical exponential distribution described by Eq. 2 (see Methods; Figure S2*B*–*D*). The mean dwell time was 61.0 s for pre-existing rodlets vs. 75.6 s for de novo rodlets; step time was 29.7 s vs. 27.3 s; and step size was 26.5 nm vs. 19.2 nm. The step rate (elongation rate within a single elongation phase, calculated as step size / step duration) was 9.0 ± 0.3 nm/min for pre-existing rodlets and 8.8 ± 0.5 nm/min for de novo rodlets. No statistically significant differences were observed for any of these parameters (Table S1). These results indicate that the elongation behavior of pre-existing and de novo rodlets did not differ significantly.

### Rodlet elongation from both ends

RolA rodlets actively elongated from both ends (Supporting Video 2). First, we defined the fast and slow ends according to the apparent elongation rate. Some rodlets in dense regions already had one side in contact with a neighboring rodlet, preventing further elongation; such rodlets were excluded from the analysis. The apparent elongation rate was 12.6 ± 2.6 nm/min for fast ends and 5.2 ± 1.2 nm/min for slow ends; the difference was statistically significant (Figure 2*A*, Table S2). Because elongation rates varied considerably among individual rodlets (Figure 2*B*), simply averaging the values at the fast and slow ends could overestimate the difference between them. Therefore, we used a linear mixed effects (LME) model (26) for statistical analysis, treating the difference between the two ends as a fixed effect and the variability among individual rodlets as a random effect. We found a statistically significant difference (*P* < 0.01) in the apparent elongation rate between the fast and slow ends. Even after accounting for variability among rodlets, this difference persisted, suggesting that it was not an artifact introduced during the analysis, but rather resulted from intrinsic differences between the two ends.

**Figure 2.**
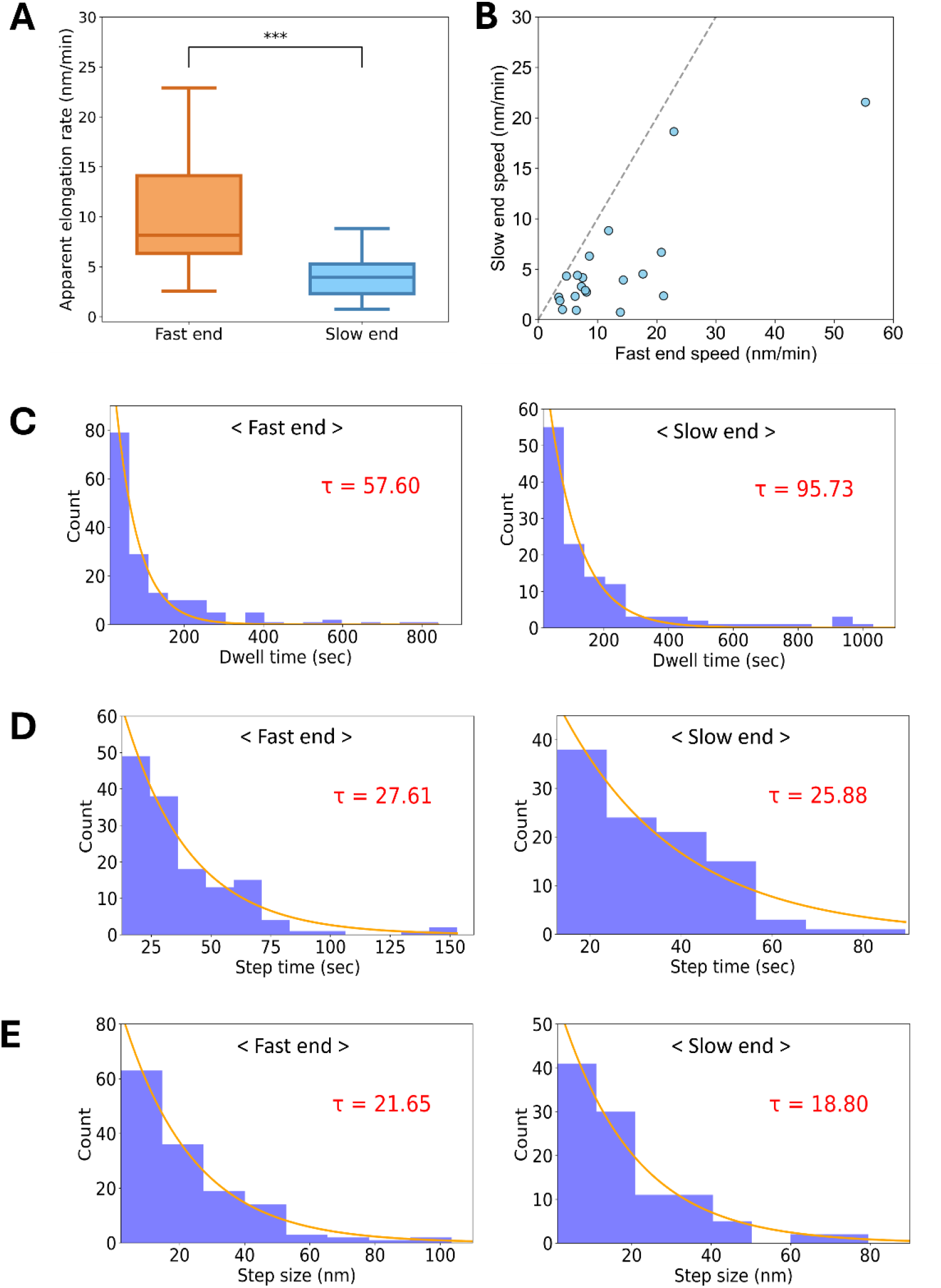
Elongation kinetics of fast and slow ends of rodlets. (A) Apparent elongation rate. Boxes extend from the 25th to 75th percentiles. The line in each box indicates the median. Whiskers reach out to the most distant point that’s still within 1.5 times the interquartile range. Statistical significance was evaluated using the Brunner–Munzel test (\*\*\**P* < 0.001). (B) Relationship between elongation rates of the fast and slow ends of the same rodlet. The dashed line is the line of equality. (C–E) Distributions of dwell time (C) and step time (D), and single-step size (E), with exponential fits (lines) giving the mean values of τ shown in each panel and Table S2.

Although rodlet elongation was polar, the mean relative difference in apparent elongation rate [(Fast – Slow) / Fast] was only about 58%. Dwell time, step time, and step size for both fast and slow ends followed the typical exponential distribution described by Eq. 2 (Figure 2*C*–*E*). The mean dwell time was 57.6 s for fast ends vs. 95.7 s for slow ends; step time was 27.6 s vs. 25,9 s; step size was 21.7 nm vs. 18.8 nm; and mean step rate was 9.2 ± 0.4 nm/min vs. 8.5 ± 0.5 nm/min (Table S2). Although no statistically significant differences were found for these four parameters (Table S2), the dwell time tended to be shorter at the fast ends than at the slow ends.

### Promotion of elongation in bundled rodlets

Multiple rodlets interacted laterally to form bundles, and individual rodlets elongated in coordination with others within the bundle (Figure 3, Supporting Video 3). No dissociation of bundled rodlets into individual rodlets was observed. Because lateral interactions between rodlets may influence their rate of elongation, we measured the elongation rates of both bundled and single rodlets; all elongating ends were included in the kinetic evaluation. The apparent elongation rate was significantly higher in bundled rodlets (13.9 ± 2.2 nm/min) than in single rodlets (7.4 ± 0.8 nm/min) (Figure 4*A*, Table S3).

**Figure 3.**
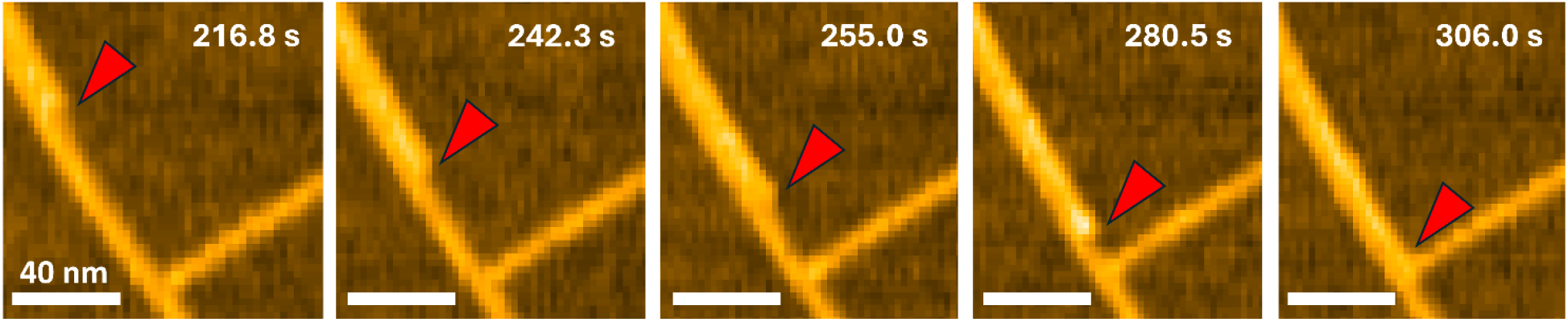
High-speed atomic force microscopy imaging of rodlet bundling. Representative images of a rodlet elongating along another rodlet. Red arrowheads indicate the tip of the rodlet during elongation.

**Figure 4.**
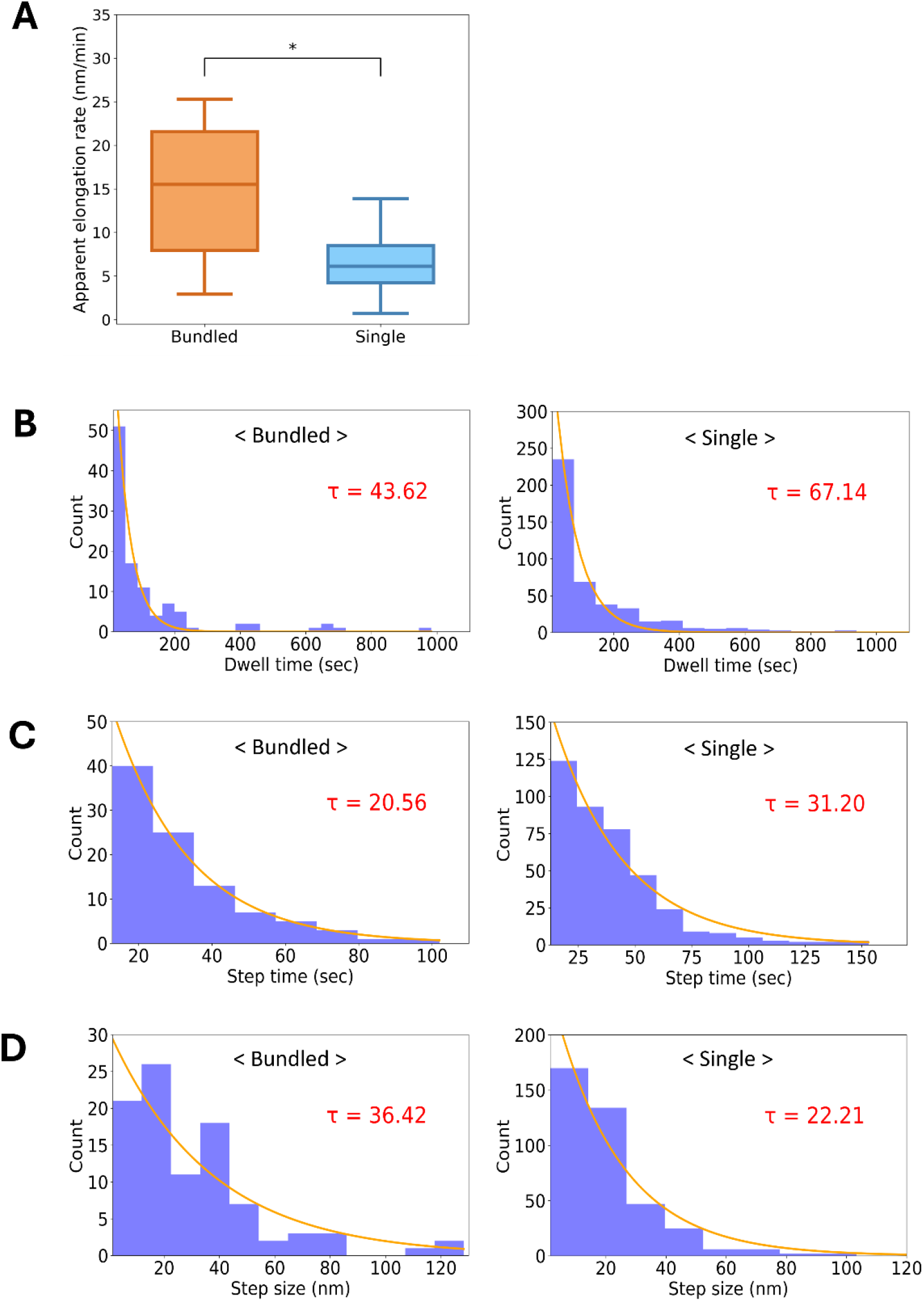
Elongation kinetics of bundled and single rodlets. (A) Apparent elongation rate. Boxes extend from the 25th to 75th percentiles. The line in each box indicates the median. Whiskers reach out to the most distant point that’s still within 1.5 times the interquartile range. Statistical significance was evaluated using the Brunner–Munzel test (\**P* < 0.05). (B–D) Distributions of dwell time (B), step time (C), and single-step size (D), with exponential fits (lines) giving the mean values of τ shown in each panel and Table S3.

In bundled rodlets, elongation and pause phases alternated and all data followed a characteristic exponential distribution described by Eq. 2 (Figure 4*B–D*). The mean dwell time was 43.6 s for bundled rodlets vs. 67.1 s for single rodlets; step duration was 20.6 s vs. 31.2 s; step size was 36.4 nm vs. 22.2 nm; and mean step rate was 17.9 ± 2.0 nm/min vs. 10.4 ± 0.6 nm/min (Figure 4*B–D*, Table S3); all these differences were statistically significant. These results suggest that the termini of bundled rodlets incorporate monomers significantly more efficiently than those of single rodlets.

### Simulations of domain formation

Monte Carlo simulations were carried out on a two-dimensional lattice to reproduce rodlet bundling and the creation of domain structures observed in nature (Figure 5). In our simulations, rodlets appeared to form domain structures when the lateral interactions between them were taken into account (Figure 5*A* and Supporting Video 4). When lateral interactions were not defined, rodlet assemblies appeared to be dispersed (Figure 5*B* and Supporting Video 5). To quantify orientation of rodlets, the angle pair correlation function was calculated. Strong lateral interactions yielded larger, more ordered domains, while weakening these interactions led to smaller domains with reduced alignment (Figure S3*A*). Under the conditions of strong lateral interactions that yielded well-ordered domain structures, the number of surface-catalyzed elongation and independent elongation of single rodlets events were counted, revealing that surface-catalyzed elongation occurred more frequently than independent elongation (Figure S3*B*)

**Figure 5.**
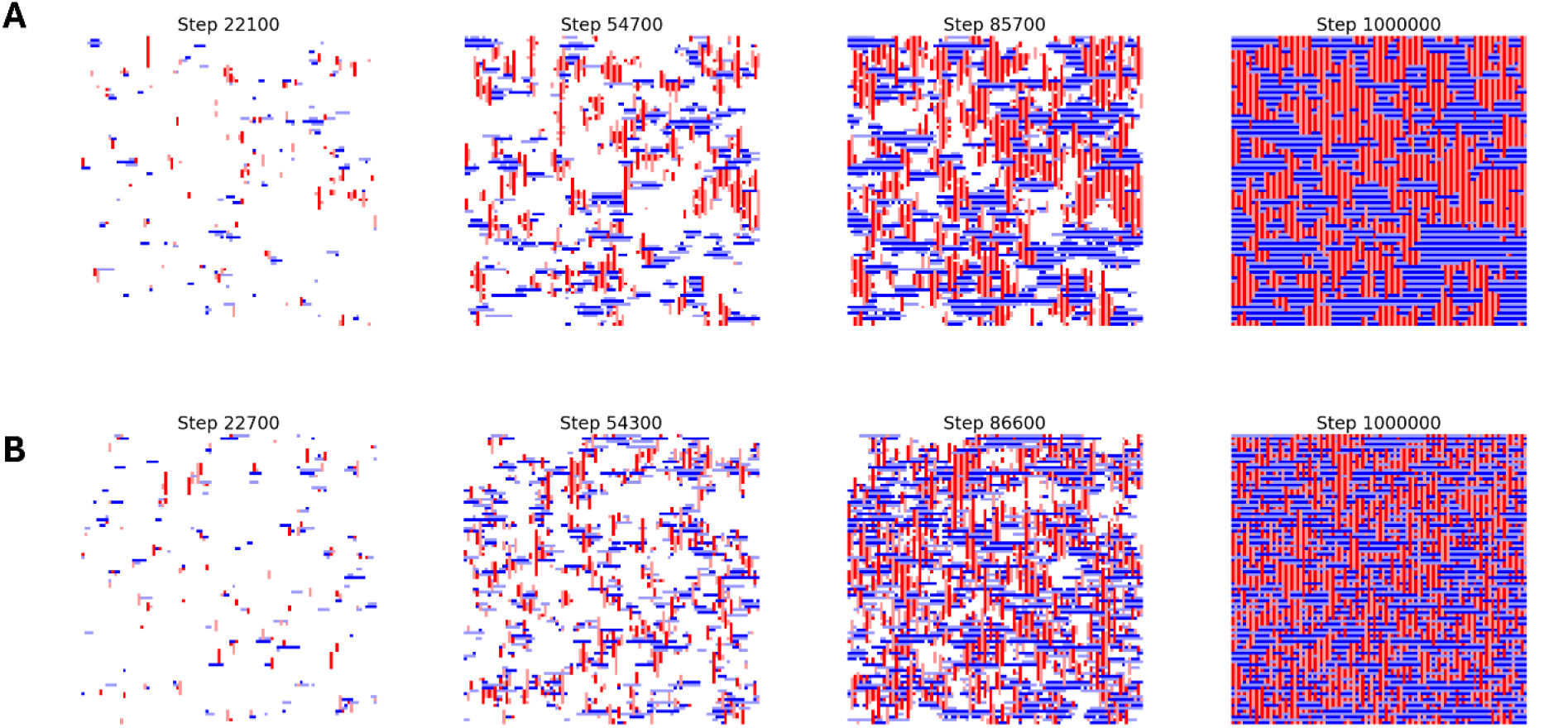
Time-lapse snapshots from Monte Carlo simulations illustrating domain formation as ε_lateral_ varies: (A) ε_lateral_ = -0.5 (lateral rodlet interactions included), (B) ε_lateral_ = 0.0 (rodlets interact independently)

## Discussion

In this study we performed bulk analysis and single-rodlet observations using HS-AFM to elucidate the dynamics of rodlet formation. CD spectra revealed that, similar to other class I hydrophobins, RolA undergoes a conformational change during self-assembly of monomers into rodlets composed of β-sheet structures (Figure S1*A* and *B*) (21, 27). In the ThT assay, no lag phase in the time course data was detected (Figure S1*C* and *D*). This result is consistent with that of Pham et al. (14) for the class I hydrophobin MPG1; in that study, the authors discussed that the rapid progression of nucleation prevented the detection of a lag phase. The reaction curve for RolA (Figure S1*C*) is also likely due to a rapid nucleation step. Although the ThT assay allows a kinetic model analysis of the reaction curves, it only provides the apparent reaction rate of the entire system, limiting the scope of the analysis and preventing a detailed investigation of the specific structural changes of RolA and the dynamics of rodlet formation.

Using HS-AFM, we monitored rodlet formation at the single-fibril level in real time and analyzed its conformational dynamics and kinetics. Our observations revealed that RolA spontaneously assembles on the substrate and transitions into rodlets, which further elongate (Figure 1*A* and *C*). Our results provide direct evidence that the nucleation–elongation model accurately describes the polymerization pathway of RolA. HS-AFM analysis also revealed that rodlet ends alternate between growth and pause states (Figure 1*D* and *E*). This suggests that the rodlet ends adopt two distinct conformational states: one that allows monomer binding and rodlet elongation and another one that prevents monomer binding, as suggested previously (28, 29). This stepwise mechanism is likely to be universal for amyloid fibril elongation, because it has also been observed in functional amyloids involved in bacterial biofilm development and in pathogenic amyloid formation (28, 30).

Our HS-AFM observations also revealed that rodlets have low polarity in the direction of extension (Figure 2*A, B*). In general, amyloid fibrils have strong polarity in their direction of extension. For example, HS-AFM and fluorescence microscopy studies of fibrils of well-characterized amyloids such as Aβ, α-synuclein, and Sup35 have shown that they elongate unidirectionally from one end or that elongation rates differ significantly between the two ends (28, 31-36). Highly polarized elongation has also been observed by HS-AFM for the secreted functional amyloid CsgA, which is involved in bacterial biofilm formation (30). Our HS-AFM analysis revealed that RolA rodlets had some degree of polarity, but elongated from both ends (Figure 1*D, E* and Figure 2*A, B*). Bidirectional elongation may provide an advantage in rodlet film formation by allowing a more efficient increase in rodlet mass from a limited number of nuclei, particularly in environments with low nucleation rates.

One of the most notable findings of this study is that the elongation rate of bundled rodlets was approximately twice that of individual rodlets (Figure 4*A*), suggesting that interactions between adjacent rodlets promote elongation. This increase in elongation rate is likely due to changes in the local environment at the rodlet ends caused by bundling. Mean dwell and step times were shorter and mean step size and step rate were greater in bundled rodlets than in single rodlets (Figure 4*B*–*D*, Table S3). The equilibrium constant was estimated as K_d_=k_open_/k_close_, where k_open_=1/τ_dwell time_, k_close_=1/τ_step time_; the K_d_ was 2.12 for bundled rodlets and 2.15 for single rodlets, indicating no significant difference between the two conditions. This suggests that the difference in chemical potential between the growth and pause states remains approximately constant regardless of whether the rodlets are single or bundled. However, because the frequency of rodlet end switching between pause and growth states is significantly higher in bundled rodlets (Figure 4*A-C*), the energy barrier for this reaction is likely lower in bundled rodlets (Figure 6). These results are consistent with the basic concept of catalysis, where the reaction rate increases without changes in the chemical potential of the reactants and products, so the results suggest that rodlets act as catalysts for the structural change of adjacent rodlet ends (Figure 6). However, this alone does not explain the higher elongation rate in bundled than in single rodlets. Fitting the data from the ThT assay indicated that RolA elongates in a dock–lock-like manner and that the locking process is the rate-limiting step in the whole reaction (Figure S1*D*), so rodlet elongation has to be considered as a multistep reaction: [Pause state] ↔ [Growth state] + [Monomer] ↔ [Growth state + Intermediate] → [Elongated rodlet]. In this scenario, the locking process ([Growth state + Intermediate] → [Elongated rodlet]) is also considered to be promoted when rodlets are bundled. The increase in step rate in bundled rodlets (Table S3) suggests a decrease in the energy barrier for the locking process. The rodlet surface can be considered to act as a reaction field that catalyzes both the structural equilibrium of rodlet ends (Growth ↔ Pause) and the locking process, increasing the elongation rate in bundled rodlets (Figure 4*A*). Three explanations are possible for the molecular mechanism. (i) A structural change at the rodlet ends promotes both structural equilibrium and the locking process. Molecular dynamics simulations have suggested that structural fluctuations at the ends of amyloid fibrils influence fibril elongation (37). (ii) RolA that undergoes docking but is not locked may interact with neighboring rodlets, thereby influencing elongation (38). (iii) RolA monomers could be attached to and concentrated on the rodlet surface, increasing the probability of their interaction with the ends of the adjacent rodlet (docking) (39). In contrast, the locking process is independent of monomer concentration and is therefore unlikely to be subject to this effect (14), but the reduction in pause state time in bundled rodlets may be explained by such monomer concentration effects. This concentration effect is easier to understand if amyloid fibril elongation is considered as one-dimensional crystal growth. In crystallization, monomers bind to the crystal surface (terrace), diffuse, and incorporate into the crystal at kink sites. In a model for amyloid fibril growth, monomers bound to the fibril surface diffuse along the lateral surface and are incorporated into the fibril ends (39). The structures of aligned fibrils of different lengths can be regarded as an arrangement where the lateral surface of long rodlets corresponds to a terrace and the ends of the bundled short rodlets correspond to kink sites. Adsorption of monomers to the lateral surface of rodlets results in a higher concentration than that in the bulk solution (40), thereby facilitating subsequent rodlet elongation, similar to one-dimensional crystal growth. Although the detailed molecular mechanisms remain unclear, it is clear that rodlets play a catalytic role in both the structural equilibrium of neighboring rodlet ends and elongation (Figure 6). Therefore, we suggest to call RolA rodlet formation pathway “surface-catalyzed elongation” and this concept is crucial for understanding the dynamics of hierarchical structure formation.

**Figure 6.**
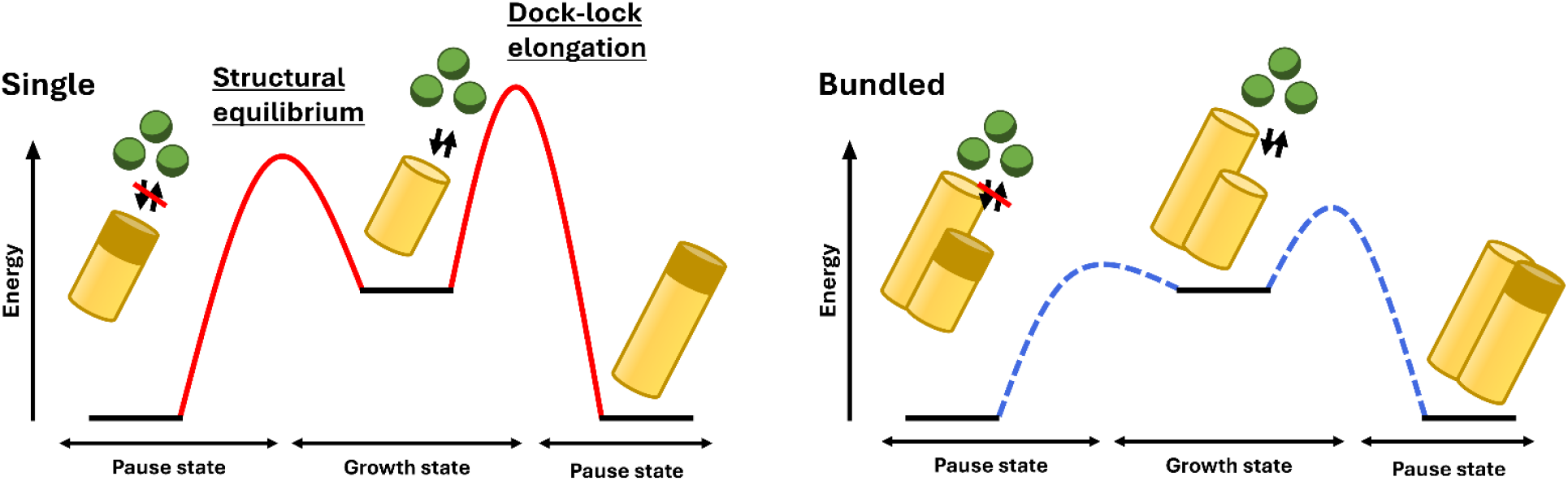
Schematic model of the stepwise growth of single and bundled rodlets. Rodlet ends have two distinct structural states: a growth state, in which monomers can bind and rodlets can elongate; and a pause state, in which monomers cannot bind. Both states are in equilibrium. When rodlets are bundled, the energy barrier between the states decreases. Rodlets elongate in a dock–lock manner, and the energy barrier of this reaction also decreases when rodlets are bundled. These decreases in energy barriers contribute to the increase in the elongation rate of bundled rodlets.

Surface catalysis in amyloid aggregation involves secondary nucleation (41), in which nucleation on the fibril surface is strongly promoted; the reverse reaction, oligomer dissociation, is also facilitated, allowing the fibril to act as a catalyst (42). Surface-catalyzed elongation differs from secondary nucleation because it promotes rodlet elongation rather than the formation of oligomers. Secondary nucleation results in a rapid increase in total fibril mass. Surface-catalyzed elongation may have a limited effect on total fibril mass because formed rodlets remain tightly packed as bundles and previously active catalytic sites become buried, preventing a net increase in their number. Surface-catalyzed elongation likely regulates fibril alignment, contributing to the formation of hierarchical structures such as large amyloid aggregates and dense ordered amyloid films. To support this hypothesis, we performed Monte Carlo simulations to reproduce rodlet bundling and subsequent domain structure formation. In our simulations, rodlets formed domain structures under conditions of strong lateral interactions (Figure 5*A*), and we observed rodlets elongated by using neighboring rodlets as observed in HS-AFM (Supporting Video 4). The appearance of our simulated structures was similar to the result of coarse-grained simulation performed by Zykwinska et al. (43). These suggest that our simulations successfully reproduce the long-timescale rodlet film formation process that HS-AFM was unable to fully capture. In addition, further detailed analysis of these simulated results quantitatively demonstrated that lateral interactions of rodlets such as surface-catalyzed elongation promote both domain enlargement and enhanced alignment (Figure S3*A*), strengthening the hypothesis that the local phenomena observed by HS-AFM can broadly influence the overall rodlet film formation. On the surface of conidia, hydrophobins typically form a tightly packed rodlet film in which rodlets are aligned in bundles (5, 6), and their appearance is similar to the results of our simulations (Figure 5*A*). Therefore, lateral interactions between rodlets observed by HS-AFM in this study may be crucial in the formation of rodlet hierarchical structures and could contribute to the morphogenesis of filamentous fungi. Lateral interactions between rodlets are likely to be mediated by specific amino acid residues of individual hydrophobin molecules exposed on the rodlet surface. However, the precise three-dimensional structure of the entire rodlet assembly remains elusive. Further structural analysis is required to elucidate the molecular mechanisms underlying surface-catalyzed elongation.

Fibril bundling has also been observed in pathogenic amyloids. Lateral interactions between fibrils of α-synuclein have been qualitatively observed using HS-AFM (31), and lateral fibril growth templated by pre-existing fibrils of Aβ42 has been qualitatively demonstrated using gold nanoparticle labeling (44). However, the kinetics and quantitative aspects of these processes have not been analyzed yet experimentally. Coarse-grained simulation has suggested that lateral interactions contribute to fibril elongation (38). Thus, although lateral interactions among fibrils and fibril bundling appear to be ubiquitous phenomena, detailed quantitative characterization of the physicochemical mechanisms at the level of individual fibrils is still lacking. The surface-catalyzed elongation observed and validated in this study may serve as a useful concept to explain the physicochemical aspects of amyloid fibril bundling.

In this study we used HS-AFM to analyze the rodlet formation dynamics of hydrophobin RolA at the single-fibril level, revealing several intriguing phenomena: (i) rodlet ends adopt two distinct structural states: a growth state and a pause state; (ii) rodlets elongate bidirectionally while maintaining polarity, although the growth rate difference between the two ends is minimal; and, the most important finding of this study, (iii) that a previously unrecognized pathway, “surface-catalyzed elongation,” governs the structural equilibrium at the rodlet ends and elongation. Because rodlets interact laterally, we propose that surface-catalyzed elongation enhances the efficiency of their assembly under high-density conditions. This work represents both an advance in understanding the cell wall surface morphogenesis of filamentous fungi by rodlets and a conceptual advance in understanding amyloid assembly, and its universality sheds light on the broader process of amyloid formation in general.

## Materials and Methods

### Purification of RolA

A strain of *A. oryzae* overexpressing wild-type RolA was created in our previous study (13) and was grown in YPM liquid medium (1% yeast extract, 2% polypeptone, and 2% maltose). In this strain, the *rolA* gene is under the control of the p-enoA142 promoter (45), which is strongly induced by maltose. Wild-type RolA was purified according to Terauchi et al. (17) as follows.

Conidia were inoculated into YPM liquid medium at 1×10^6^ conidia/ml and cultivated at 30 °C for 48 h with shaking (160 rpm). The culture broth was filtered through Miracloth (Merck KGaA, Darmstadt, Germany) to separate the mycelia from the culture supernatant. The supernatant was adjusted to pH 8.5 using 0.1 M Tris solution (pH 10.5), Milli-Q water was added to adjust the electrical conductivity to 1.0 mS/cm, and the solution was applied to a Cellufine Q-500 column (Seikagaku Co., Tokyo, Japan) equilibrated with 5 mM Tris-HCl buffer (pH 9.0). RolA was eluted with a 0–0.3 M linear gradient of NaCl. Each fraction was checked by sodium dodecyl sulfate– polyacrylamide gel electrophoresis (SDS-PAGE; 3% stacking gel and 17.5% running gel). The fraction containing RolA (13.6 kDa) was dialyzed against 10 mM sodium citrate buffer (pH 4.0) and applied to an SP-Sepharose Fast Flow column (GE Healthcare, Chicago, IL, USA) equilibrated with the same sodium citrate buffer. RolA was eluted with a 0.05–0.3 M linear gradient of NaCl and the fractions (3 ml each) were collected into test tubes containing 1 ml of 100 mM Tris to neutralize. Each fraction was checked by SDS-PAGE, and the fraction containing RolA was dialyzed against 10 mM ammonium acetate (pH 7.0) and lyophilized. All the purification procedures were performed at 4 °C. Lyophilized RolA was dissolved in Milli-Q water or buffer before use.

### Measurements of CD spectra

RolA monomer solution (50 µg/ml) was prepared in 10 mM sodium acetate buffer (pH 5). Rodlet solution was prepared by vortexing the monomer solution for 1 h at room temperature (23 °C). Each solution (200 µl) was dispensed into a 1-mm quartz cuvette (GL Sciences, Tokyo, Japan). The spectra were measured in a J-725 CD spectrometer (JASCO International, Tokyo, Japan) over a wavelength range of 190–250 nm at 23 °C with a bandwidth of 1 nm and step intervals of 1 nm. The sample compartment was continuously flushed with N_2_ gas. The spectra were averaged over 5 scans. The baseline was recorded using 10 mM sodium acetate buffer (pH 5).

### Thioflavin T assay

Because ThT specifically binds to rodlets and emits strong fluorescence (14, 18), it allows evaluation of the rodlet formation process. Lyophilized RolA was dissolved in 10 mM sodium acetate buffer (pH 5); ThT dissolved in sodium-acetate buffer, sodium acetate buffer, and RolA solution were added to the wells of a black 96-well microplate (Nunclon Delta Surface; Thermo Electron Corporation, Waltham, MA, USA) in this order to a final ThT concentration of 20 µM, and fluorescence was measured (430 nm excitation/480 nm emission) in a microplate reader (Fluoroskan Ascent, Thermo) with shaking at 600 rpm (shaking diameter = 3 mm) at 30 °C. The model in Eq. 1 below was used to interpret the obtained curves (14):

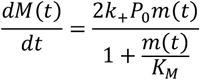

where *M*(*t*) is the total rodlet mass, *m*(*t*) is the monomer mass, *P*_0_ is a constant (number of open ends), *k*_+_ is the elongation rate constant, and *K*_*M*_ is the Michaelis constant of elongation, which gives the monomer concentration at which the effect of saturation becomes important. This model, proposed by Pham et al. (14), assumes that elongation proceeds in two steps, with the monomer first “docking” at the rodlet end, followed by structural reorganization (“locking”) to form a stable structure, suggesting that monomer concentration-independent locking becomes rate-limiting when the monomer concentration is sufficiently high.

### Preparation of hydrophobically modified SiO_**2**_ substrate

The silicon substrate surface was hydrophobically modified according to Terauchi et al. (17). First, silicon wafers (p-Si wafers, ≤0.02 Ω; Mitsubishi Material Trading Co. Ltd., Tokyo, Japan) were consecutively ultrasonically cleaned for 15 min each with chloroform, acetone, and 2-propanol to remove microscopic contaminants from the surface. Then, the wafers were cleaned with an ultraviolet ozone cleaner (SKB401Y-01; Sunenergy Co. Ltd., Kanagawa, Japan) for 30 min each on the back and surface to remove organic contamination on the surface, immersed in chloroform solution containing 0.1% *n*-octyltrichlorosilane (Tokyo Chemical Industry Co., Ltd., Tokyo, Japan) and left overnight at room temperature (23 °C) to modify the surface. The surface was then dried with N_2_ gas, and the wafers were stored in a desiccator.

### HS-AFM imaging

A custom-built HS-AFM instrument (20, 46) equipped with a small cantilever (BL-AC10DS-A2; Olympus, Tokyo, Japan) was operated in tapping mode with the cantilever oscillating at its resonance frequency in liquid (∼0.4 MHz) and an amplitude of ∼2.0 nm (free oscillation amplitude ∼2.5 nm) at room temperature (23 °C). The probes were prepared on the top of cantilevers by electron beam deposition in a field emission scanning electron microscope. HS-AFM scanned 2 × 2 µm of the observation area with 500 × 250 pixels at 12.75 s per frame. Samples were prepared as follows. Rodlet formation on the surface of a droplet was induced by incubating 20 µl RolA solution (3.68 µM) in 10 mM sodium acetate (pH 5.0) on Parafilm for 5 min at room temperature (23 °C). The droplet surface was then pressed against a hydrophobically treated SiO_2_ substrate for 30 s, which caused rodlet transfer to the substrate. The substrate with the attached rodlets was placed in the sample chamber of the HS-AFM instrument filled with 10 mM sodium acetate (pH 5.0). The size and shape of the rodlets were determined by HS-AFM and the buffer in the chamber was replaced with 7.35 µM RolA solution (100 μg/ml) in the same buffer. The choice of RolA concentration was based on the results of the ThT assay (Figure S1*D*) (18), which indicated that at 100 μg/ml RolA its concentration is not a rate-limiting factor for rodlet elongation, whereas the locking process is rate-limiting independent of monomer concentration. Then, rodlet elongation was imaged by HS-AFM for up to 30 min. HS-AFM movies (image sequences) were processed using ImageJ 1.53e software (https://imagej.nih.gov/ij/) according to Watanabe-Nakayama and Ono (47). From the HS-AFM videos that captured rodlet elongation, the coordinates of all individual rodlet ends were measured by using ImageJ to obtain the time history of their positions. The rodlets were divided into the following categories for analysis: (i) those that were already attached to the substrate at the start of the observation period (pre-existing); (ii) those newly formed from monomeric RolA in the chamber that became attached to the stage during the observation period (de novo); (iii) those that were alone on the stage (single); and (iv) those located next to each other (bundled). In cases where elongation stopped due to a rodlet collision, we analyzed only the data recorded immediately before the collision. The numbers of rodlets used for measurements are presented in Tables S1–S3. From these time courses, dwell time, step time, and step size were determined and statistically analyzed. Dwell time, step time and step size had typical exponential distributions (Figure 2*C-E*, 4*B-D*, S2*B-D*), as expressed by Eq. 2:

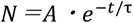

where *N* is the number of events, *A* is a constant, *t* is the dwell or step time, and *τ* is the mean time.

### Monte Carlo simulations

Monte Carlo simulations on a two-dimensional lattice were performed to model the formation of the hierarchical structure of rodlet assemblies based on previous study and with modifications (43). Particles representing short rodlets were placed on the cells and allowed to interact with rodlets existing in the neighboring cells. Each cell could take one of three states: empty (0) or containing a vertical or a horizontal rodlet (−1, 1). The simulation started with all lattices empty. At each simulation step, one of the empty cells (*i, j*) in the lattice was randomly selected and a rodlet was placed into this cell (corresponding to nucleation). If the four neighboring cells were empty, the orientation of the newly placed rodlet was selected randomly (−1 or 1 with probability 1/2 each). The frequency of this random nucleation is governed by the probability *P*, which in this case was set at 0.005. If some rodlets are present in the neighboring cells, the orientation of the newly placed rodlet is determined by their interaction energy (defined below) with the neighboring rodlets. The vector of the newly placed rodlet (*i, j*) is *v* and the vector of its neighboring rodlet (*i’, j’*) is *w*. The joint vector between the newly placed rodlet and its neighbor is *b*=(*j*−*j’, i*−*i’*). If the newly placed rodlet is oriented in the same direction as the neighboring rodlet, then |*v*·*b*|=1, and if the two rodlets have different orientations, then |*v*·*b*|=0. When the orientation of the newly placed rodlet aligned with that of the neighboring rodlet (*v*=*w*) and it was placed in the extension direction (|*v*·*b*|=1), the rodlet was considered to have elongated, and the depth of potential energy between them was defined as ε_elongation_. If *v*=*w* but the new rodlet was placed laterally (|*v*·*b*|=0), the depth potential energy between them was defined as ε_lateral_. If the orientation of the new rodlet did not align with that of the existing rodlet (*v≠w*), the depth of potential energy was set to 0. Surface-catalyzed elongation gave ε_elongation_ +ε_lateral_. The frequency of nucleation when neighboring rodlets were present was set to 1.0 if the newly placed rodlet was aligned with the extension direction, and 0.1 otherwise. Simulations were performed with the following parameters: lattice size, 100×100; number of simulation steps, 1,000,000; temperature, *k*_B_*T*=0.1, where *k*_B_ is the Boltzmann constant. Six simulations were performed with ε_elongation_ fixed at −1.0 and ε_lateral_ varied over six values: 0.0, −0.01, −0.1, −0.2, −0.5, −1.0. The time evolution of the Monte Carlo simulations was governed by the Metropolis algorithm (48). To evaluate the degree of rodlet orientation, we calculated the angle pair correlation function (49,50). The size and degree of alignment of a domain structure can be evaluated by examining how well the direction of rods correlates with that of distant rods. The angle pair correlation function is defined in following Eq. 3:

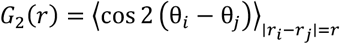

where θ_i_ and θ_j_ are the angles of rods *i* and *j*, respectively, and |*r*_i_ −*r*_j_|=*r* is the Euclidean distance between them. *G*_2_(*r*)=1 indicates an ordered structure in which all rods are perfectly oriented, whereas *G*_2_(*r*)=0 indicates a disordered structure with randomly oriented rods.

## Acknowledgments

This work was supported by the Japan Society for the Promotion of Science under a Grant-in-Aid for Scientific Research (B) (23K26810) to K.A. and a Grant-in-Aid for JSPS Fellows (24KJ0419) to N.T., and by the Noda Institute for Scientific Research (K.A.) and Bio-SPM Collaborative Research Proposal (K.A.) from the WPI-Nano Life Science Institute, Kanazawa University and World Premier International Research Center Initiative (WPI), MEXT, Japan.

