## Supporting Information for "High-speed atomic force microscopy reveals surface-catalyzed elongation mechanism of fungal functional amyloid, hydrophobin RolA"

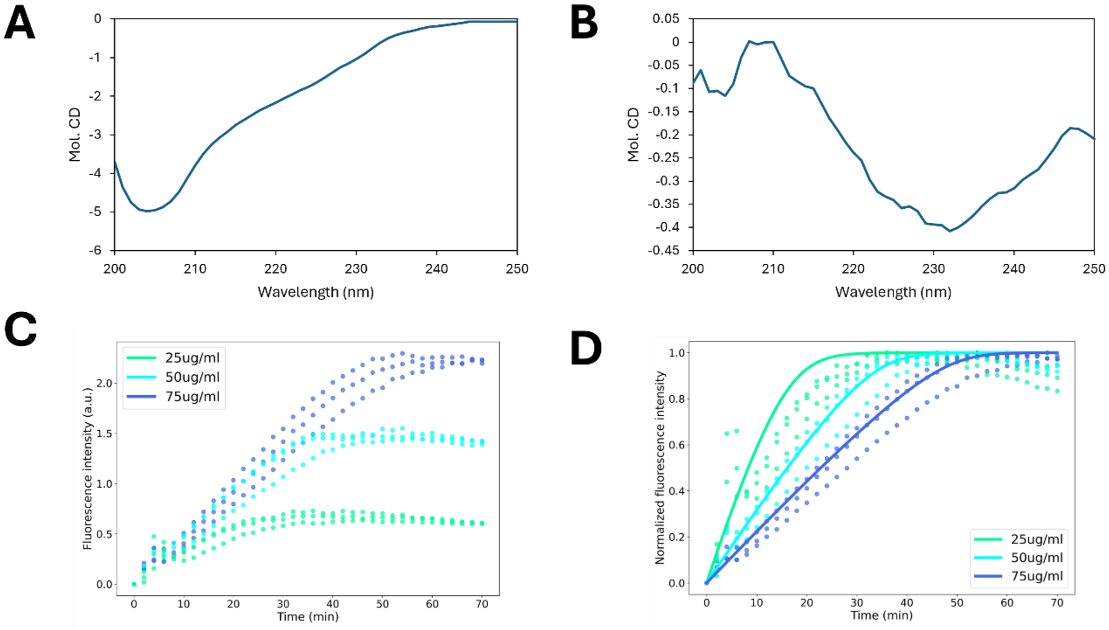


**Fig. S1.** Formation of RolA rodlets in bulk solution. (A, B) CD spectra of 50 μg/ml RolA in the monomeric state (A) and rodlet state (B), indicating a conformational change through self-assembly. (C, D) Time course of ThT fluorescence at different initial RolA concentrations at 30 °C with shaking for raw data (C) and normalized data (D). Dotted lines, measured data; solid lines, the results of fitting using Eq. 1 (see Methods).

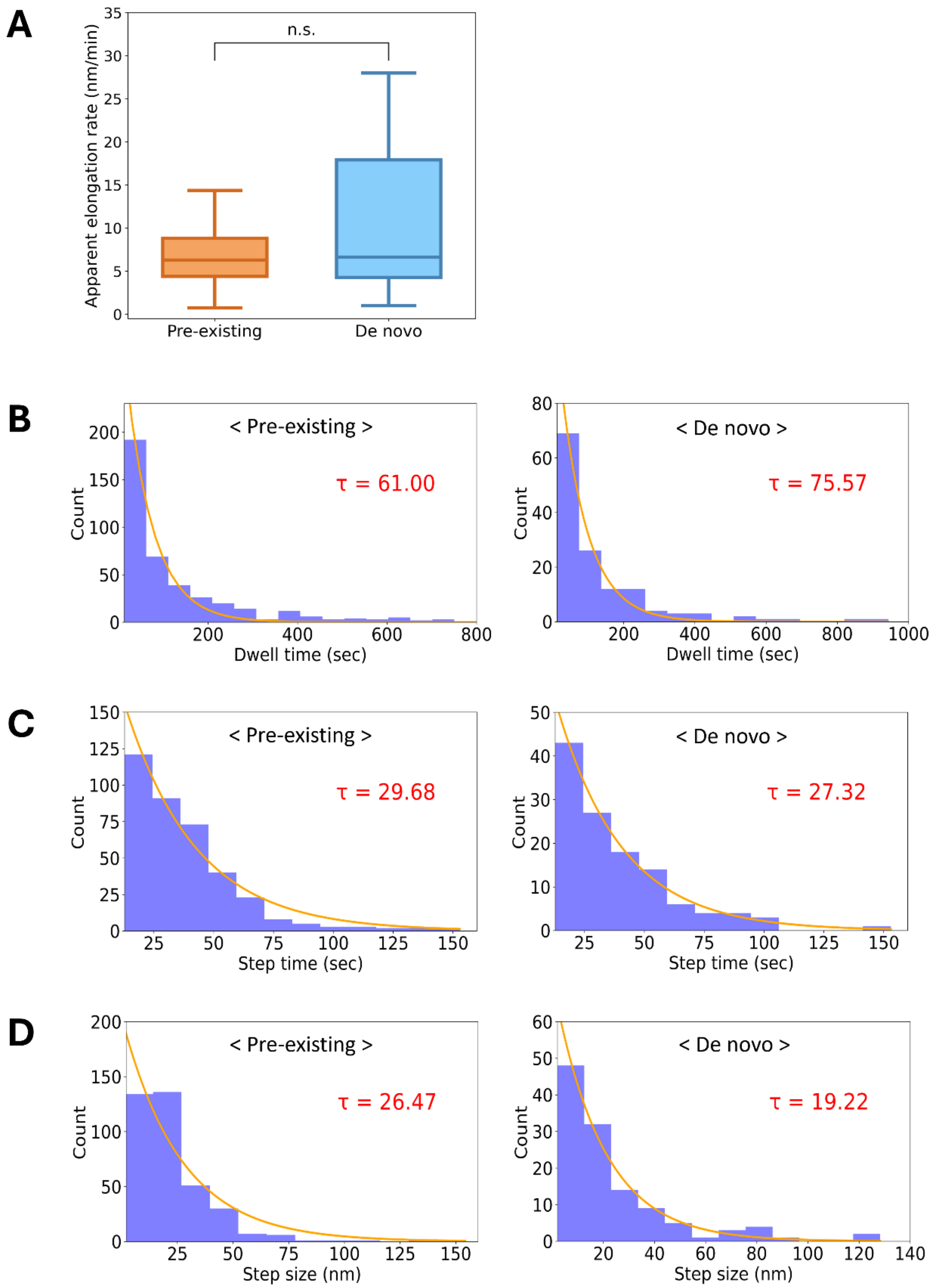


**Fig. S2.** Elongation kinetics of pre-existing and de novo rodlets. (A) Apparent elongation rates. Boxes extend from the 25th to 75th percentiles. The line in each box indicates the median. Whiskers reach out to the most distant point that's still within 1.5 times the interquartile range. Statistical significance was evaluated using the Brunner–Munzel test; n.s., not significant. (B–D) Distribution of dwell time (B), step time (C), and single-step size (D), with exponential fits (lines) giving mean values of τ shown in each panel and Table 1.

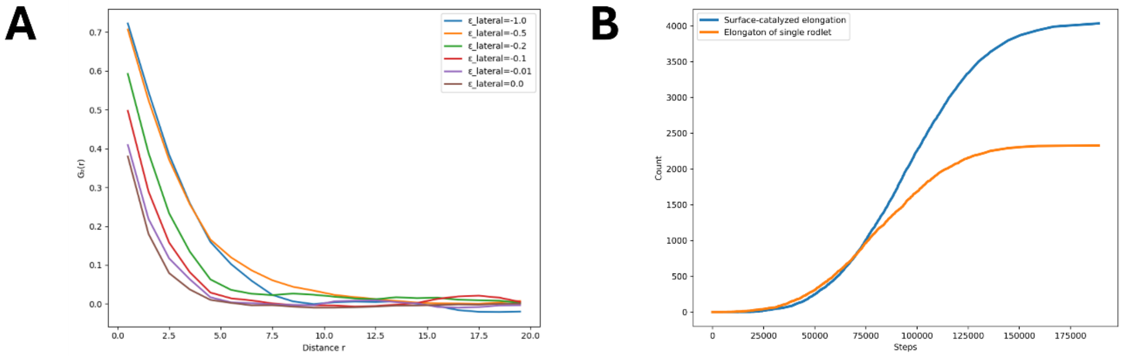


**Fig. S3.** Results of Monte Carlo simulations of the effect of lateral rodlet interactions on domain formation. (A) Contribution of surface-catalyzed elongation to domain formation. (B) Angle pair correlation function in different ε_lateral values.

**Table S1.** Kinetic parameters of pre-existing and de novo rodlets.

| | Pre-existing | De novo | *P* value* |
|---|---|---|---|
| Number of analyzed rodlets | 51 | 24 | – |
| Total number of analyzed steps | 411 | 137 | – |
| Apparent elongation rate (nm/min) | 7.9 | 11.6 | 0.609 |
| Mean dwell time (s) | 61.0 | 75.6 | 0.747 |
| Mean step time (s) | 29.7 | 27.3 | 0.872 |
| Mean step size (nm) | 26.5 | 19.2 | 0.250 |
| Step rate (nm/min) | 9.0 | 8.8 | 0.462 |

* *P* values were calculated by the Brunner–Munzel test.

**Table S2.** Kinetic parameters of fast and slow ends.

| | Fast | Slow | *P* value* |
|---|---|---|---|
| Number of analyzed rodlets | 20 | 20 | – |
| Total number of analyzed steps | 158 | 126 | – |
| Apparent elongation rate (nm/min) | 12.6 | 5.2 | $8.33 \times 10^{-5}$ |
| Mean dwell time (s) | 57.6 | 95.7 | 0.0556 |
| Mean step time (s) | 27.6 | 25.9 | 0.432 |
| Mean step size (nm) | 21.7 | 18.8 | 0.0682 |
| Step rate (nm/min) | 9.2 | 8.5 | 0.0906 |

* *P* values were calculated by the Brunner–Munzel test.

**Table S3.** Kinetic parameters of bundled and single rodlets.

| | Bundled | Single | *P* value* |
|---|---|---|---|
| Number of analyzed rodlets | 13 | 62 | – |
| Total number of analyzed steps | 105 | 441 | – |
| Apparent elongation rate (nm/min) | 13.9 | 7.4 | 0.0140 |
| Mean dwell time (s) | 43.6 | 67.1 | 0.0244 |
| Mean step time (s) | 20.6 | 31.2 | 0.0131 |
| Mean step size (nm) | 36.4 | 22.2 | $1.53 \times 10^{-4}$ |
| Step rate (nm/min) | 17.9 | 10.4 | $1.31 \times 10^{-7}$ |

* *P* values were calculated by the Brunner–Munzel test.

**Movie S1** (separate file).
HS-AFM video of entire field of view. HS-AFM scanned 2 × 2 μm of the observation area at 12.75 s per frame. The video is played back at ×50 higher speed.

**Movie S2** (separate file).
HS-AFM video of rodlet elongation from both ends. Enlarged video of 2 × 2 μm scan video at 12.75 s per frame. The video is played back at ×50 higher speed.

**Movie S3** (separate file).
HS-AFM video of rodlet bundling. Enlarged video of 2 × 2 μm scan video at 12.75 s per frame. The video is played back at ×50 higher speed.

**Movie S4** (separate file).
Time-lapse video of Monte Carlo simulation considering lateral interactions between rodlets. $\varepsilon_{elongation}$, -1.0; $\varepsilon_{lateral}$, -0.5; lattice size, 100×100; number of simulation steps, 1,000,000; $k_BT$=0.1.

**Movie S5** (separate file).
Time-lapse video of Monte Carlo simulation assuming no lateral interactions between rodlets. $\varepsilon_{elongation}$, -1.0; $\varepsilon_{lateral}$, 0.0; lattice size, 100×100; number of simulation steps, 1,000,000; $k_BT$=0.1.
